# Prospects and Challenges for Graphene Drums As Sensors Of Individual Bacteria

**DOI:** 10.1101/2023.11.20.567863

**Authors:** I.E. Rosłoń, A. Japaridze, L. Naarden, L. Smeets, C. Dekker, A. van Belkum, P.G. Steeneken, F. Alijani

## Abstract

Graphene-drum-enabled nanomotion detection can play an important role in probing life at the nanoscale. By combining micro- and nanomechanical systems with optics, nanomotion sensors bridge the gap between mechanics and cellular biophysics. They have allowed investigation of processes involved in metabolism, growth, and structural organisation of a large variety of microorganisms, ranging from yeasts to bacterial cells. Using graphene drums, these processes can now be resolved at the single-cell level. In this perspective, we discuss the key achievements of nanomotion spectroscopy, and peek forward into the prospects for application of this single-cell technology in clinical settings. Furthermore, we discuss the steps required for implementation and look into applications beyond microbial sensing.

## Introduction

Since the discovery of cells by Robert Hooke and Antoni van Leeuwenhoek, mankind’s understanding of biological systems at the cellular level has kept pace with the progress in microscopic tools to observe and study such systems. It is not surprising that advancements such as fluorescence microscopy (*1*) and cryogenic electron-microscopy (*2*) were pivotal in the development of cellular biology. Yet, the observation of many cellular processes *in vivo* remains a significant challenge (*3*), sincecertain processes in cells cannot be easily visualized due to their small signal amplitudes and high levels of noise.

In this light, the recent realization that even single-cellular organisms generate small mechanical fluctuations with a broad spectrum of frequencies, might be viewed as a next step in our technical advancement of studying cellular processes. Longo and colleagues (*4*) did Atomic Force Microscope (AFM) cantilever experiments that revealed that populations of living bacterial cells (100-1,000 cells) generate nanomotion on cantilever sensors. Inspired by these experiments, we developed the tools to use graphene drums as sensors (*5*) that are capable of recording the “beating” of even individual bacteria. Our nanomotion sensors encompass an ultra-thin suspended two-dimensional (2D) graphene membrane with relatively low stiffness (*k* = 0.1 N/m) that is sensitive enough to transduce forces as small as a picoNewtons - even in the oxygenated liquid environment that is required to keep the microbial cells alive. We showed that single-bacteria emit small nanometer-scale vibrations when alive, that can be recorded by these nanomechanical sensors. Such vibrations may provide insight into the metabolic activity and processes taking place inside a single cell.

In this perspective, we first highlight the scientific achievements of the nanomotion techniques, especially when applied to single cells. We then address the prospects for application of single-cell nanomotion technology in clinical settings, where it can enable Rapid Antibiotic Susceptibility Testing (RAST), where we demonstrate single-cell nanomotion signals from clinical isolates of five different bacterial species. Next, we describe the challenges in performing sensitive and specific high-throughput graphene based RAST and discuss application of alternative read-out techniques and materials for single-cell nanomotion sensors. Finally, we summarize the wide range of possibilities to use this technology in various fields beyond bacterial sensing, ranging from probing fundamental biophysical processes to yeast activity monitoring and protein force sensing.

### 1 Recent advances in nanomotion spectroscopy

Nanomotion spectroscopy consists of attaching micro-organisms to a mechanical structure and measuring the nanoscale vibrations that the organism induces (*6*). Cantilever sensors have been first used to detect the nanomotion of groups of bacteria, but also of various other cells, such as yeasts and other eukaryotes (*7*). The technique has attracted particular interest for screening of slowly growing pathogens (*8*). The cantilever is moved by the forces produced by the live specimen (Figure 1a), and the deflection is recorded via the reflection of a laser on a 4-quadrant photo diode or through coupling with fibre optics (*9*).

**Figure 1:**
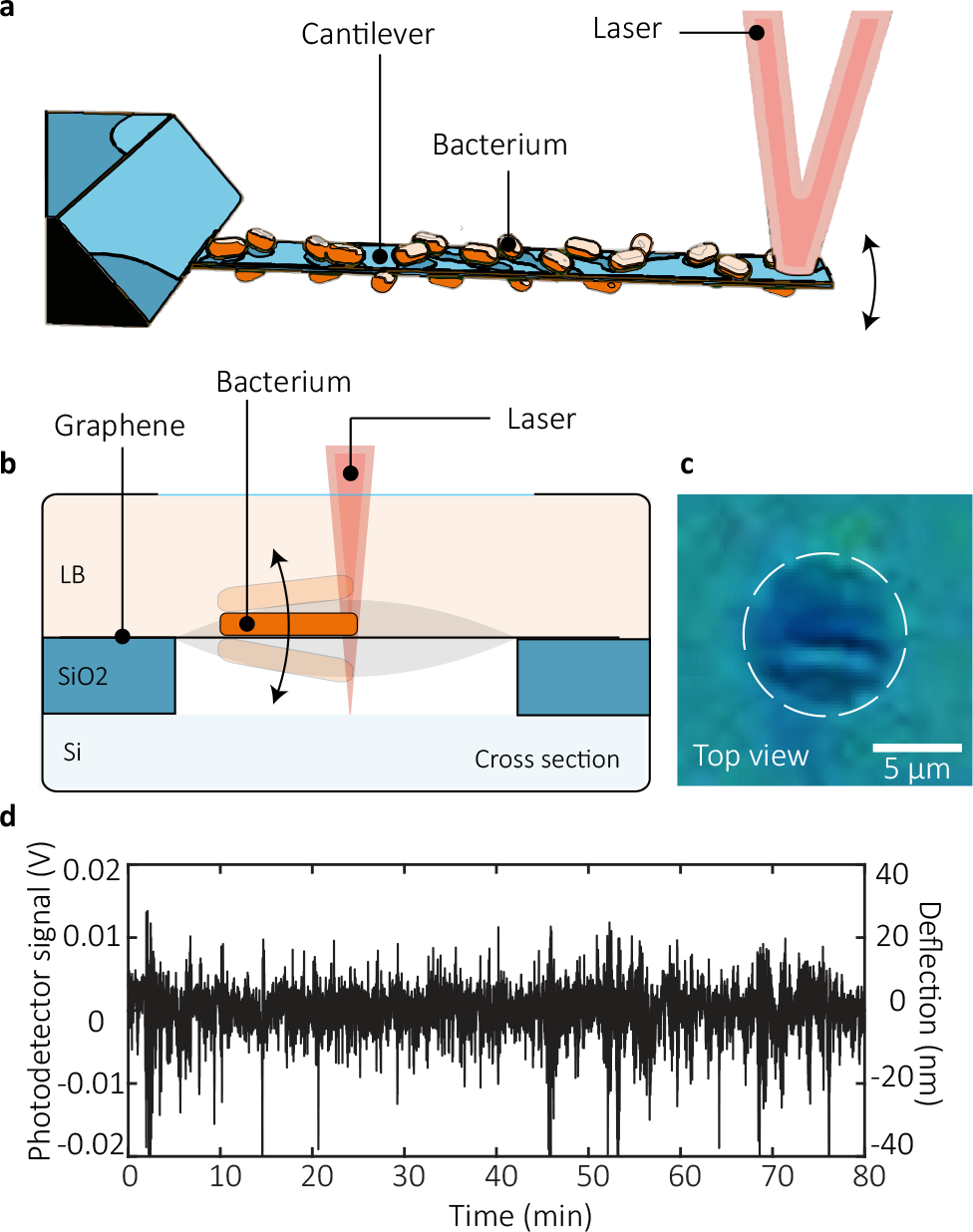
(a) Nanomotion of bacterial cells was first detected by measuring groups of cells adhered to cantilever sensors; figure adapted from (*11*). (b) With grapheme drums, it is possible to observe the nanomotion of even single bacterial cells. (c) An optical image of a single bacterial cell attached to a graphene drum. The drum is outlined by a dashed white circle. (d) Nanomotion signal obtained by measuring a single cell under constant conditions for more than an hour. Large oscillations occur with an amplitude of up to 40 nm. Figures adapted from (*5*).

Recently, we introduced a new method for probing nanomotion of single bacteria. By using suspended graphene drums (*5*), the mechanical time-amplitude data trace of a single bacterium adhered to the drum can be obtained using laser interferometry (Figure 1b,c). In this technique, the vibrations induced by the single bacterium moves the mechanical receiver which in turn changes the optical characteristics of the cavity underneath the graphene. As a result, the drum displacement can be read-out optically by measuring the intensity of the reflected light. Schematics of both setups can be seen in Figure 1 alongside a typical drum deflection trace caused by a single bacterium. In approximation, the deflection *δx* depends on the force *F* exerted by the bacterium on the flexible support, *F≈ kδx*, where the out-of-plane stiffness *k* of the flexible support determines the sensitivity.

#### 1.1 Graphene drums

Graphene drums have interesting properties that make them excellent candidates for the role of flexible support for nanomotion-enabled activity detection. They are ultra-thin, are virtually mass-less, have very low stiffness but at the same time have high tensile strength which prevents them from breaking under tension from the liquid environment (*10*). Important further aspects for this kind of detection method are threefold: first, the size of the sensitive area needs to match the object of interest. By matching the size of the detector to that of the specimen, effects of background environmental signals can be minimized. The displacement detector also needs to be highly compliant (i.e. have low mechanical stiffness). A low stiffness allows the detector to be easily moved by any external impetus, therefore increasing the minimal detectable force. Finally, appropriate optical properties are required in order to translate the microbial motion effectively into a readable signal. A perfect device for nanomotion detection combines these characteristics in the most efficient manner. For these reasons, graphene sensors are an ideal candidate to play the role of flexible support for nanomotion detection.

#### 1.2 Data acquisition and analysis

The characteristic sensing signal is a noisy intensity trace versus time, where the information of interest is contained in the noise fluctuations. The typical approach for analysing nanomotion time data consists of two steps. The first is a drift subtraction, which is done by subtracting a linear fit from the raw data, over a range of several seconds to minutes. Subsequently, the variance *σ*^2^ is most commonly used as metric (*11, 12*), although more elaborate metrics have also been conceived (*6*). In nanomotion-based bacterial motility and viability testing, the change in variance is generally expressed with respect to a control sample. That means, that changes and differences in nanomotion are compared to a reference value of the variance exhibited by this control sample.

Cantilevers are generally covered by large numbers of bacteria, such that the recorded signal is enough to be detected. Typical ensembles are 100 to 1000 cells. This makes it difficult to discern specific signals from single cells, but does provide an average representation of an entire population. This may allow for the detection, in real-time, of bacterial variants such as persisting cells. On the other hand, by obtaining a distinctive signal from single cells, one not only can start using nanomotion for identification and analysis of mixtures, but it also allows to look deeper into the cellular mechanisms that cause this variance. Two studies also performed Fast Fourier Transform (FFT) analysis of the cellular signals and found a 1*/f* ^*α*^ type of signal, which is common among biological samples (*5, 13*). Despite the apparent similarity of the signals at first sight, it is worth exploring to see if Artificial Intelligence (AI) or more intricate signal-analysis techniques can distinguish or identify different cells solely by the emitted nanomotion.

### 2 Road to applications in clinical settings

Accurate identification and Antibiotic Susceptibility Testing (AST) of bacteria is crucial for clinical microbiology laboratories to guide appropriate treatment and infection control. However, culture-based AST methods, which are commonly used, are time-consuming, require one or more days to identify resistant pathogens and even longer to provide antibiotic susceptibility profiles (*14*). In parallel, incubation periods in blood culture systems commonly range from 1 to 3 days (*15*). An additional challenge is that some pathogenic bacteria are fastidious, which means that are difficult to impossible to grow in laboratory conditions because they have complex or restricted nutritional and environmental requirements, such as bacteria from the *Legionella* or the *Bartonella* genera (*16*). As a result, broad-spectrum antibiotics are often administered to patients, while physicians still await AST results. The implementation of faster broad-spectrum AST technologies will have a large impact on clinical outcomes (*17*). This is because early and precise differential diagnosis of infections is critical for reducing morbidity and mortality of patients, hence reducing healthcare costs (*18*). In the long-term, this will lead to a societal benefit of reduced development of antibiotic resistance by making sure the right antibiotic is given for the correct duration and with the right formulation.

Applications of nanomotion spectroscopy as RAST sensors in clinical settings (*19*) is of great interest, and might even lead to simultaneous identification and susceptibility testing of bacteria, reducing the time and resources required for the overall testing process. Most importantly, a key challenge lies in the robustness and throughput levels of such a technique before it can be widely introduced in clinical practice.

#### 2.1 Emerging industrial platforms

Current wide-scale operating platforms such as the bioMérieux Vitek 2 and the BD Phoenix already generate relatively rapid results (typically in 10-18h), but require a standardized microbial sample, which still requires culturing of the specimen for 24-48h and identification of the pathogen (*20*). There are multiple platforms under development for rapid AST technologies, with time to result below 6 hours (*21*). Optical detection platforms, such as Gradientech and BacteriScan, are the most similar to the widespread systems already in use in clinical practice (*22, 23*). These platforms optically determine turbidity changes of the incubated sample and generate AST results within 3 hours. However, these new platforms do not have the ability to perform simultaneous identification or test directly on non-urine samples, let alone perform tests on single cells.

Another branch of emerging technologies bases its rapid AST on bacterial DNA extraction and subsequent genomic testing, such as the platforms of GenomeKey and Day Zero Diagnostics (*24, 25*). This approach is suitable for simultaneous identification and susceptibility testing at an increased throughput, and might offer a way to work directly with non-purified specimens if sufficient sensitivity is achieved. Genomic techniques rely on a library of DNA sequences encoding the resistance, which need to be known upfront to allow for detection of a resistance. However, genes are not necessarily expressed, which might lead to disagreements between genomic and culture AST results.

Nanomotion spectroscopy techniques are under development by SoundCell and Resistell, from which the latter is currently conducting clinical test in a tertiary-care hospital (*26*). Resistell is developing a cantilever-based nanomotion method whereas SoundCell bases the readout on graphene drums. These technologies might provide rapid AST within 2 hours as well as simultaneous identification and susceptibility testing, but increasing throughput would require microfluidics accommodating multiple cantilevers or arrays of drums.

#### 2.2 Trials on clinical strains

Here, we discuss the applicability of the graphene RAST on different classes of clinically relevant bacterial strains. We performed measurements on isolates of *Escherichia coli, Klebsiella pneumoniae, Methicillin-resistant Staphylococcus aureus (MRSA), Streptococcus agalactiae* and *Pseudomonas aeruginosa* see Fig.2. With this selection, we covered species that have the highest frequency of incidence and high prevalence of both infection and antibiotic resistances (*27*). These species covered both gram negatives (*E. coli, P. aeruginosa, K. pneumoniae*) and gram positives (MRSA, *S. agalactiae*), as well as motile and non-motile strains. For each species, we confirmed the presence of nanomechanical fluctuations (Figure 2). The obtained nanomotion signals were processed following the procedure outlined above and discussed in (*5*) which involves calculating the variance *σ*^2^ of the signal, or its motion amplitude *σ*, which is a measure of bacterial viability before and after adding an antibiotic. A high value of *σ* means the bacteria are metabolically active and alive, while a value close to baseline, due to nanomotion of the suspended graphene alone (see 2c), means they are not. In Figure 2d, we also show the difference in the power spectral density (PSD) of a drum with and without bacteria, which are clearly different.

**Figure 2:**
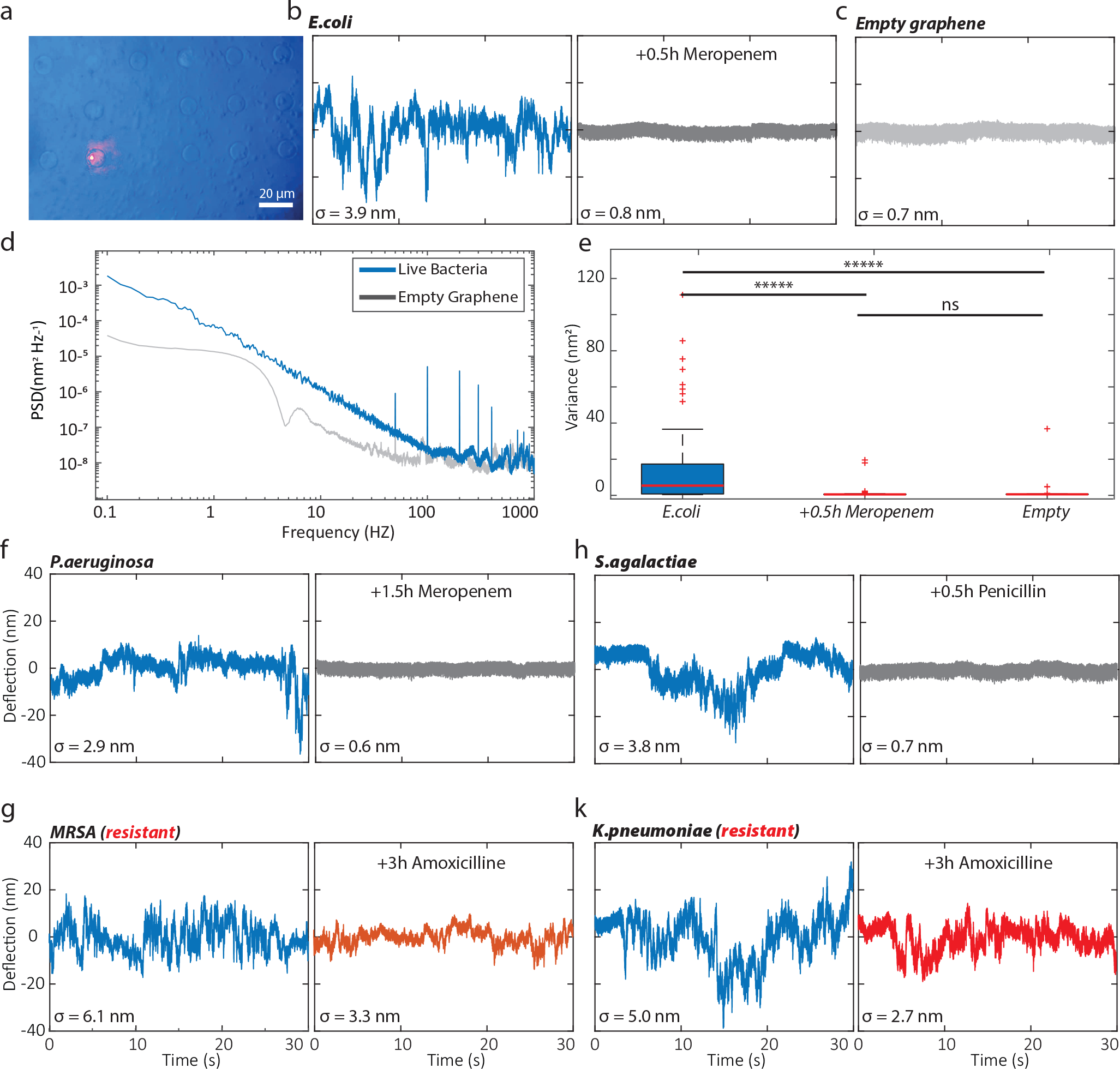
Experiments on clinical isolates of various bacterial species. (a) Snapshot image of a RAST sample showing graphene drums in the presence of bacteria; (b) Nanomotion signal from clinical isolates of *E*.*coli* before and after adding meropenem; (c) Nanomotion signal of an empty graphene drum ; (d) Power spectral density (PSD) of a graphene drum with and without bacteria; (e) Box plot showing the statistical analysis performed on a number of graphene drums with and without antibiotics. The statistical data are collected from 80 measurements (ns stands for not significantly different); (f) Nanomotion signal of clinical isolates of *P. aeruginosa* before and after adding meropenem; (h) Nanomotion signal of clinical isolates of *S. agalactiae* before and after adding Penicilin; (g) Nanomotion signal of *MRSA* before and after adding Amoxicillin; (h) Nanomotion signal of *K*.*pneumoniae* before and after adding Amoxicillin. For each species a 30s initial trace is shown in blue, followed by a trace in gray (susceptible) or red (resistant) recorded after administering a bactericidal concentration of the antibiotic. Meropenem was used at a final concentration of 1 *µ*g/mL, Penicillin at 0,125 *µ*g/mL, and Amoxicillin at 60 *µ*g/mL. The value of motion amplitude *σ* is shown next to each trace.

In all cases we added various antibiotics close to the antibiotic breakpoint concentrations, a defined threshold concentration of an antibiotic that helps categorize bacterial isolates into susceptible and resistant categories (*28*), (1 *µ*g Meropenem at 1 *µ*g/mL for *E. coli* and *P*.*auruginosa*, Penicillin at 0,125 *µ*g/mL for *S*.*agalactiae*, and Amoxicillin at 60 *µ*g/mL for *MRSA* and *K*.*pneumoniae* (both highly resistant strains). After administering the antibiotics we re-measured the same cells after just half an hour to two hours of incubation. For cases where the strains were susceptible, even within half an hour a significant signal drop was observed (see Figure 2b-h). The signals recorded on susceptible cells after adding antibiotics were indistinguishable from that of an empty graphene drum, indicating that these antibiotics were indeed effective in killing the cells. Importantly, when the experiments were performed with resistant strains (see Figure 2g,k) no significant changes were observed. Even after 3 hours of exposure to the drug, the cells still displayed nanomotion significantly higher than the background signal.

Single-cell diagnostics hold the promise of unprecedented precision and rapid turnaround time (*29*), and in this respect graphene-based nanomotion RAST has a particularly good potential as it may work on samples from clinical isolates, and yields results within mere hours. However, the current graphene RAST technology requires highly skilled personnel for the preparation of the clinical samples, and only one sample at a time can be tested in the pilot setup. Furthermore, the trial was performed on clinical isolates with prior identification of species. The development of a complete RAST platform, thus, requires realization of further steps in terms of high-throughput and lowered manual labour demand from operating personnel.

### 3 Outlook and directions for development

Multiple technical developments of the technique are foreseen in this section, including upscaling with microfluidics, nanomotion-pattern-based cell recognition, and alternative read-out techniques for realizing highthroughput sensing. At a more fundamental level, all the root causes of nanomotion have not been untangled yet, although there is a clear indication that flagellar activity contributes significantly to single bacterium nanomotion(*5*). Possible further mechanisms that can be held accountable for nanomotion signals are also looked into in this section.

#### 3.1 Parallel read-out with high speed and high throughput

To bring single-cell nanomotion spectroscopy into clinical practice, the first step is to enhance the throughput. The read-out, for instance, can be enhanced by engineering a detection methodology for rapid detection of many graphene drums in parallel. Measuring cells one-by-one is a time consuming process and especially for screening purposes it is highly recommended to parallelize the process (*30*). This challenge can be tackled by recording the signal from several drums simultaneously, either by a “scanning” over a set of drum positions, or by illuminating multiple drums and recording intensity data at once with multiple detectors (or a camera with sufficiently high frame rate). Scanning over a set of drums allows the use of a photodetector with high dynamic range, whereas the camera approach allows for massively parallelized measurements at the expense of dynamic range. Also automated read-out cartridges could greatly simplify usage of the technology, and put lower demands on operating personnel, in turn realizing higher throughput and accuracy. Such read-out cartridges could accommodate multiple sensor chips to simultaneously test different antibiotics, at various concentrations to determine microbiologically relevant metrics such as Minimum Inhibitory Concentration (MIC). Ultimately, a measurement system might be expanded in size and throughput to screen multiple cartridges in one session, or conversely shrunk in size to be used as a random access diagnostics tool.

#### 3.2 Machine Learning for cell identification

It is of great interest to examine whether the nanomotion signals from the drums can be used to identify various bacterial species. For instance, differentiating between gram negative and gram positive species, such as *E*.*coli* and *S. aureus* in a fast and reliable manner would be of high importance for clinicians. This may now be achieved thanks to single-cell information that graphene nanomotion sensors do obtain. In order to perform this, the use of Artificial Intelligence (AI) algorithms would be ideal. However, for such an approach to be effective, the algorithm must be first trained on a considerable amount of nanomotion data for different types of bacterial species and strains thereof. A potential scheme to realize this vision and to identify if a sample contains resistant or susceptible bacteria, is provided in Figure 3a. Once the AI algorithm is trained on a large library of samples including empty drums, different classes of bacteria as well as resistant and susceptible strains, first, a sanity check is performed to recognize if the drum is suspended and suitable for measurement. Then, the algorithm can make distinction between drums that are containing a bacterium or not. Measurements are only valuable when the drum is intact and contains a bacterium. Next, a distinction can be made on the type of bacteria based on the signal they emit. Finally, the control and antibiotic treated data can be compared to obtain a result for the susceptibility test.

**Figure 3:**
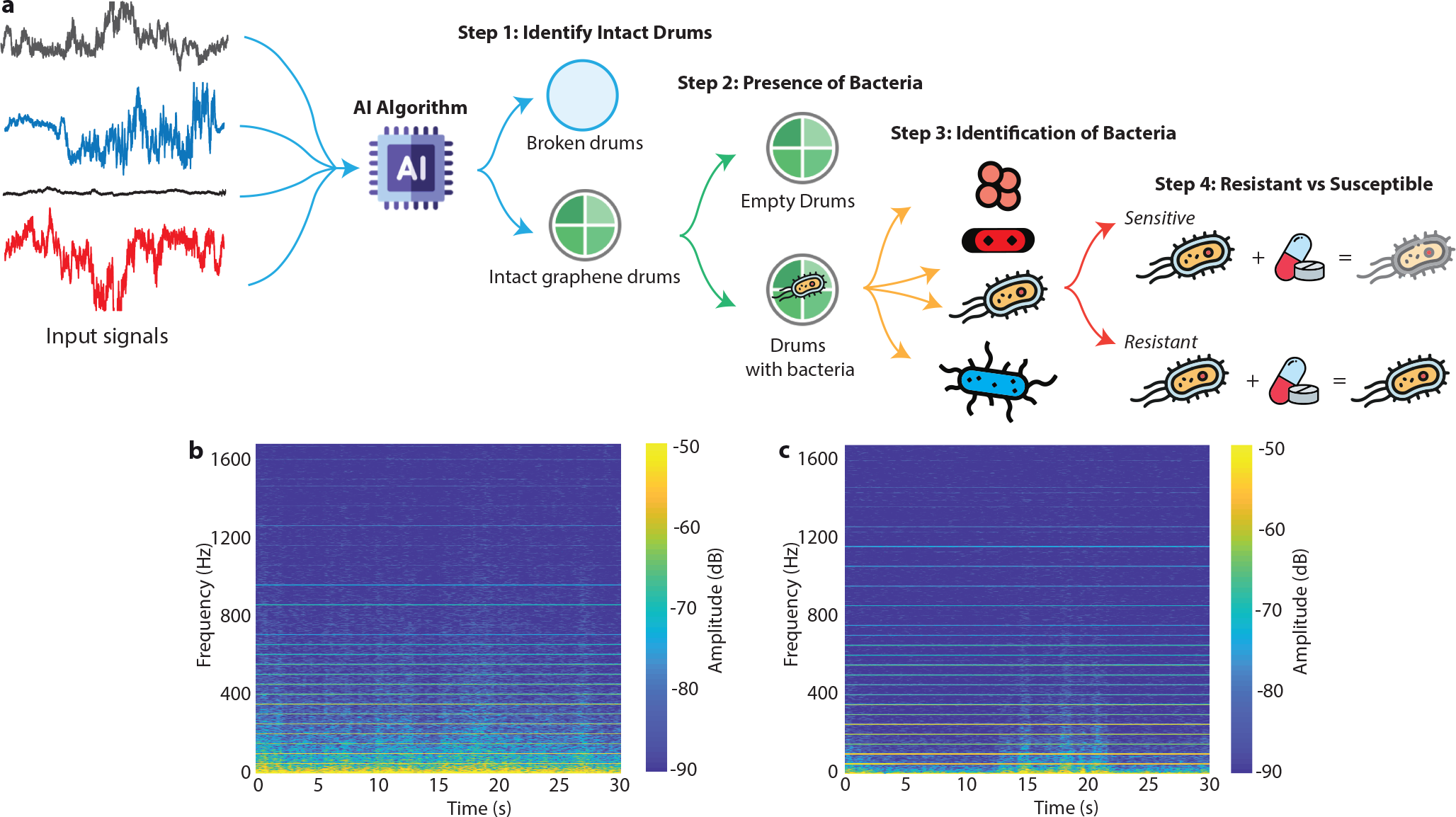
Potential process flow for combined identification and RAST based on single-cell nanomotion signals and AI algorithms. (a) To determine if an unknown bacterial sample is sensitive (susceptible) or resistant to an antibiotic, nanomotion measurements are first to be checked upon the intactness of the drum and the presence of a bacterium. After these checks are passed, the bacterial species can be classified, and finally its resistance judged. (b-c) Spectrograms obtained by applying the short-term Fourier transform to nanomotion measurement traces of a control (b) and antibiotic treated (c) bacterium. The low frequency components (*<* 200 Hz) clearly differentiates the two spectograms. Such spectograms can be used as an input for an AI algorithm to perform swift image based classification. Horizontal lines are at multiples of 50 Hz and due to mains interference.

First efforts on using Machine Learning for automated classification of susceptible and resistant strains have already been reported (*8*). Further development might benefit from pre-processing of the raw data, such as short-term Fourier transform (STFT) analysis (*31*), to limit the computational effort required for swift image based classification. Figure 3b-c show the STFT of a control and test sample and a low frequency component on the spectogram is visible as a differentiating feature. AI is well suited for the analysis of a large volume of data to recognize such patterns that might even not be readily discernible by the human eye.

The use of AI for the identification of bacteria via signals obtained from their nanomotion is motivated by several factors. There is a high automation potential that these algorithms can offer to the process of analyzing nanomotion signals and identifying bacteria, which reduces the reliance on manual labor while increasing efficiency. AI algorithms are capable of adapting to new data and improving its accuracy over time as more data are collected, making it a suitable tool for the dynamic field of bacterial identification.

#### 3.3 Alternative read-out techniques and 2D material substrates

In nanomotion experiments reported thus far, two kinds of optical read-out methods have been used to measure deflection of the mechanical lever. Either the angle under which an incoming beam is reflected can be measured, or the change in the reflectivity which causes a light intensity modulation. In both cases signals can be acquired with a photodiode or a high-speed camera. In either case the methodology requires for a laser beam to shine through the growth medium, which provokes fierce design requirements on both the measurement chamber and the microscope in terms of materials used and environmental noise suppression, rendering it harder to use outside of academic setting.

The usage of 2D material drums on silicon allows for other read-out techniques, among which specifically electronic read-out embedded on the chip is of interest. Various schemes can be considered, such as capacitive coupling to the membrane (*32*), embedded strain gauging within the suspended layer (*33*) and even integrated photonics (*34*). Such a read-out system allows the development of this technology into a self-contained lab-on-a-chip platform that includes the processing logic on-board. Such a solution would be especially interesting for point-of-care testing where simplicity and cost of use are major decisive factors (*35*). Further research could also be aimed at identifying other viable 2D materials next to graphene. So far, only silicon cantilevers, as well as bilayer graphene have been used as base material for a flexible support, yet there is plethora of different 2D materials that can be used with potential in nanomotion spectroscopy that is unknown (*36*).

#### 3.4 Cell deposition and selectivity

Manifold immobilization strategies exist for targeted attachment of living cells to a sensitive surface (*37*), which by themselves can enhance the quality and selectivity of the obtained signal, as long as the cell growth and viability are not hindered. Manipulating the surface characteristics of the graphene to make it selectively sticky to cells would be a development of great benefit. By patterning the adhesive substrate such that only the suspended areas of the graphene accept cells, it should be possible to work with smaller aliquots of bacterial samples. If the adhesive surface is also cell- or biomarker-specific, separate areas on one chip could be used for trapping different species. This may allow one to test even complicated samples such as direct patient samples typically containing a mixture of cells.

#### 3.5 Probing cellular dynamics as root causes of nanomotion

By probing the nanoscale motion, one could investigate which processes occur in single cells without intervening in them. Preliminary analyses of the nanomotion signals (*38, 39*), have suggested that flagellar activity is a main contributor to nanomotion (*5*), but the correlation between the measured signal and its physical source is not unravelled yet. Various processes, such as cell viability (*40*), osmotic pressure fluctuations, metabolic activity and organelle mechanics might all contribute, as depicted in Figure 4, in addition to the environment acting as a possibly equally important contributing factor. Active conformational changes in topoisomrases have been shown to generate nanomotion (*41*). Furthermore, in eukaryotes intracellular organelles, such as mitochondria, which are responsible for energy generation, also show nanomotion (*42*). Detecting nanomotion using fluorescent labelled products or organelles (*43*) is an interesting way to further explore nanomotion causes which might lead to new insights into the root cause.

**Figure 4:**
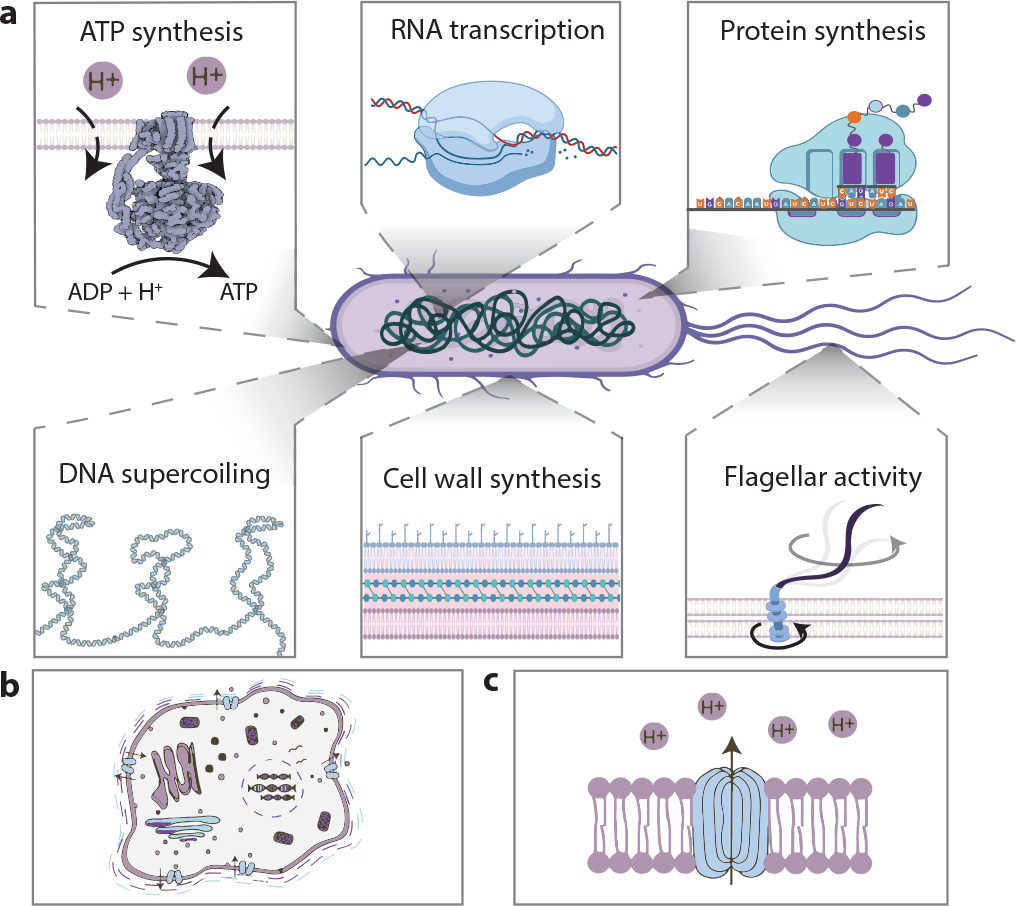
Root causes of nanomotion in bacterial cells. (a) Various processes in the bacterial cell can be responsible for the mechanical nanomotion observed, such as ATP synthesis, RNA transcription, protein synthesis, DNA supercoiling, cell wall synthesis and activity of flagella and pili. Flagellar activity has been shown to be a major contributor to the observed nanomotion. (b) Eukaryotes contain intracellular organelles that can generate nanomotion, such as mitochondria, which are responsible for energy generation. (c) Active ion channels can also generate nanomotion due to their conformational changes. Figures adapted from (*5, 39*).

### 4 Applications beyond bacterial sensing

Over the course of the past decade the nanomotion technique has been applied to various different species, and it is applied with success for bacteria, yeast, neurons, and other mammalian cells. Most of the research mentioned was performed on AFM cantilevers, rather than graphene drums, as the latter so far was only used for detection of single bacterial cells.

#### 4.1 Yeasts and bio-industrial applications

Yeasts are used in many biotechnological applications, ranging from food production chains to constituents of bioreactor flora (*44*). They play a significant role in the industrial production of biofuels and enzymes. For all these applications, it is of major interest to verify the activity and thus productivity of yeast strains before a bioreactor is populated. Early and massively parallel screening is a good strategy to alter and verify the quality of yeasts with a faster turnover, thus finding superior industrial traits earlier. The nanomotion that can be measured from yeasts, alike bacteria, is most likely directly linked with their metabolic activity (*45*). In most cases, a higher metabolic activity will translate into a higher production of the yeast’s industrially relevant compound. We envisage therefore, that by probing the nanomotion of the yeast’s, the productivity of strains can be directly measured and potentially improved.

#### 4.2 Molecular force monitoring

The high force sensitivity of graphene might enable sensing beyond the limit of single cells. Some molecules are active as a result of light (*46*) or solute concentration (*47*), and can perform mechanical work. It will be interesting to see if graphene membranes can be used as a detector for probing the forces exerted by these molecules during mechanical events, such as DNA supercoiling or protein folding. Here, a significant challenge will lie in the preparation of such samples, and the controlled attachment of the biomolecules onto the graphene surface.

### 5 Final remarks

In recent decades rapid advancements in microfabrication technology are generating new areas of application in biology. The wide availability of microelectromechanical systems (MEMS) since the 1990’s has provided researchers new platforms to experimentally study cell mechanics and their mechano-microbiology. With the development of the graphene drums as sensors for single cells, it is now possible to measure and analyze cellular dynamics even at the level of single bacteria. This raises thrilling prospects for usage of nanomotion detection for both identification as well as antibiotic susceptibility analysis. In our opinion, the development of massively parallel graphene nanomotion sensors can be a gamechanger in this field. The ability to robustly run even thousands of nanomotion spectroscopy measurements in parallel will open the way towards development of robust RAST sensors combined with nanomotion based identification.

## Methods

### Sample preparation

All experiments were performed on anonymous clinical isolates of *E*.*coli, K*.*pneumoniae, MRSA, S. agalactiae* and *P. aeruginosa* cells obtained from the medical microbiology department of the Reinier Haga Medical Centre in Delft. We grew cells in Muller-Hinton Broth overnight at 30 degrees Celsius to reach the late exponential phase. On the day of the experiment, the overnight culture was refreshed (1:100 volume) for 2.5 h in fresh broth at 37 degrees Celsius to reach an optical density (OD600) of 0.2–0.3. Then 10 ml of the refreshed culture was mixed with (3-Aminopropyl)triethoxysilane (APTES, Sigma-Aldrich) to reach a final concentration of 0.1% (volumetric). This acts as a binder between the bacteria and the chips. A chamber with a graphene-covered chip inside was then filled with the solution, which was left for 15 minutes in a horizontal position to deposit the bacteria on the surface. Afterwards, the chamber was flushed with broth to prevent additional bacteria from depositing and maintain an average coverage of a single bacterium per drum. The setup was equipped with nano positioners (Attocube ECSx5050) that allow for automated scanning over an array of drums. The motion of the bacterium was transduced on the drum and recorded using a digital oscilloscope.

### Graphene chip fabrication

Experiments are performed on circular suspended graphene membranes. A silicon wafer with a silicon dioxide layer is patterned by etching holes in the silicon dioxide, where the silicon acted as stop layer, resulting in 285 nm deep circular cavities with diameters ranging from 2 to 10 *µ*m. Graphene resonators are fabricated by suspending single and few-layer graphene over circular cavities using a dry transfer technique. Both exfoliated graphene flakes and chemical vapor deposited layers are used as resonator. The samples are annealed in an Argon furnace at 400K to remove all polymer residuals. The setup consists of a red laser aimed and focused at a Fabry-Pérot cavity formed by the bottom silicon layer and the suspended graphene layer. The deflection of the graphene layer along the optical field of the red laser modulates the reflected light intensity that can be read out by a photodiode. The setup allows detection of the absolute deflection of the membrane.

### Data processing

All data are collected and plotted using MATLAB code. For analysis, the same routines are used as described earlier in (*5*). For the short term fourier transform a custom code was written in MATLAB, with the following settings: blackman type window with a length of 2048 and an FFT length of 8192.

## Data availability

The data that support the findings of this study are available from the corresponding authors upon request.

## Acknowledgements

Financial support was provided from the European Union’s Horizon 2020 research and innovation programme under ERC starting grant ENIGMA (no. 802093, F.A. and I.E.R.), ERC PoC GRAPHFITI (no. 966720, F.A. and A.J.), Dutch Research Council (NWO) take-off grant, Graphene Flagship (grant nos. 785219 and 881603, P.G.S.). We also acknowledge financial support from European Innovation Council Transition Grant (no. 101136371, I.E.R, A.J., P.G.S. and F.A) as well as UNIIQ: Finance the Future and Graduate Entrepreneur Fund.

## Author contributions

All authors contributed to the analysis, interpretation of the results and writing of the manuscript, with a main contribution to the writing by I.E.R. R.A. performed the bacterial manipulation and experiments. I.E.R. constructed the setup and performed the interferometry experiments. The project was supervised by P.G.S. and F.A.

